# Alzheimer’s disease-associated protective variant Plcg2-P522R modulates peripheral macrophage function in a sex-dimorphic manner

**DOI:** 10.1101/2024.05.28.596275

**Authors:** Hannah A. Staley, Janna Jernigan, MacKenzie L. Bolen, Ann M. Titus, Noelle Neighbarger, Cassandra Cole, Kelly B. Menees, Rebecca L. Wallings, Malú Gámez Tansey

**Affiliations:** Department of Neuroscience, University of Florida College of Medicine, Gainesville, Florida, USA; Center for Translational Research in Neurodegenerative Disease, University of Florida College of Medicine, Gainesville, Florida, USA; McKnight Brain Institute, University of Florida, Gainesville, Florida, USA; Norman Fixel Institute for Neurological Diseases, University of Florida Health, Gainesville, Florida, USA

**Keywords:** Alzheimer’s disease, macrophage, PLCG2, sex differences

## Abstract

Genome-wide association studies have identified a protective mutation in the phospholipase C gamma 2 (*PLCG2*) gene which confers protection against Alzheimer’s disease (AD)-associated cognitive decline. Therefore, PLCG2, which is primarily expressed in immune cells, has become a target of interest for potential therapeutic intervention. The protective allele, known as *P522R*, has been shown to be hyper-morphic in microglia, increasing phagocytosis of amyloid-beta (Aβ), and increasing the release of inflammatory cytokines. However, the effect of this protective mutation on peripheral tissue-resident macrophages, and the extent to which sex modifies this effect, has yet to be assessed. Herein, we show that peripheral macrophages carrying the *P522R* mutation do indeed show functional differences compared to their wild-type (WT) counterparts, however, these alterations occur in a sex-dependent manner. In macrophages from females, the *P522R* mutation increases lysosomal protease activity, cytokine secretion, and gene expression associated with cytokine secretion and apoptosis. In contrast, in macrophages from males, the mutation causes decreased phagocytosis and lysosomal protease activity, modest increases in cytokine secretion, and induction of gene expression associated with negative regulation of the immune response. Taken together, these results suggest that the mutation may be conferring different effects dependent on sex and cell type, and highlight the importance of considering sex as a biological variable when assessing the effects of genetic variants and implications for potential immune system-targeted therapies.

## BACKGROUND

Alzheimer’s disease (AD) is a devastating neurodegenerative disorder that afflicts millions of people worldwide over two-thirds of which are women (1). Classically, AD is histologically characterized by the presence of amyloid-beta (Aβ) plaques extracellularly, and the formation of tau tangles within neurons. However, over recent years, the immune system has been implicated in the pathogenesis and progression of the disease. Of particular interest are cells of the innate immune system, which includes both peripheral monocytes and macrophages, and the brain-resident macrophages, microglia. These cells are considered to be the body’s first line of defense, and are main propagators of inflammation, which has been shown to exacerbate neurodegeneration (2–4). Furthermore, these cells, having the ability to phagocytose, can take up these proteinaceous Aβ plaques as well as dying neurons. It has been noted for some time that although they possess the ability to do so, microglia and peripheral monocytes and macrophages fail to effectively clear these plaques. This has been hypothesized to occur with declining immune function with age, because as the disease progresses, the cells of the immune system become dysregulated, senescent, and associated with low-grade chronic inflammation (5). The lack of disease-modifying interventions for AD until recently has fueled the search for identification of dysregulated immune pathways as potential new targets to delay, arrest, and perhaps prevent disease development (6).

Genome-wide association studies identified a mutation within the phospholipase C gamma 2 (*PLCG2*) gene, called *P522R*, that protects against AD-related cognitive decline (7, 8). PLCG2 is highly expressed in immune cells including brain-resident microglia, peripheral monocytes, and tissue-resident macrophages (9). Determining how this mutation confers its protection in these cell populations will allow us to better understand how these cells play a role in the progression of AD, and may open up new avenues for therapeutic development. PLCG2 has been shown to support numerous myeloid cell functions including phagocytosis, cytokine secretion, survival, and lipid metabolism (10). To do so, it acts downstream of receptors such as triggering receptor expressed on myeloid cells 2 (TREM2) and toll-like receptors (TLRs) to cleave phosphatidylinositol (4,5) bisphosphate (PIP2) into diacylglycerol (DAG) and inositol 1,4,5 triphosphate (IP3). DAG and IP3 go on to both release intracellular calcium and activate downstream pathways including nuclear factor light chain enhancer of activated B cells (NF-κB) (10, 11). The *P522R* mutation has been shown to be hypermorphic, indicating an enhancement of immune cell functions (12). As the primary brain cells that express PLCG2 are microglia, much of the work aimed at investigating how its protective effect is conferred has focused on how it alters microglial function. Studies have shown that the *P522R* mutation increases the release of inflammatory cytokines by microglia, and has variable effects on phagocytosis depending on cargo size (13). For example, studies showed that the *P522R* mutation reduced phagocytosis of *Escherichia coli* but increased phagocytosis of Aβ (14, 15). Further *in vivo* work confirmed this increased uptake of Aβ in a 5XFAD mouse model of amyloid-pathology (16).

However, the effect of this mutation in peripheral tissue-resident macrophages has yet to be studied. As this protective pathway is of interest in the development of potential therapeutics, it is important then to understand how those therapeutics may affect the immune system globally, and not just within the brain. Furthermore, peripheral macrophages may not just be bystanders in the progression of AD, as it has been shown that they are not only more efficient phagocytes of Aβ than microglia, but they are also capable of traversing the blood-brain barrier during times of injury and disease (17–21). There is further evidence to suggest this traversal of peripheral myeloid cells to the brain may not even be necessary to impact AD progression. Indeed, studies have shown that AD progression and pathology can be directly impacted by peripheral immune cell functions. For example, Aβ efflux out of the brain and into peripheral reservoirs such as the blood and tissues has been shown to occur (22, 23). The clearance of Aβ from these reservoirs was associated with lower cerebral Aβ pathology (24). Furthermore, peripherally derived Aβ has been shown to enter the brain and exacerbate neuropathology, further supporting the importance of peripheral Aβ stores and clearance (25). Therefore, determining the effect of this protective mutation on peripheral macrophage function is of vital importance.

Additionally, given that women are at higher risk for the development of AD, we posit that sex-dependent differences in the immune system may be partially responsible for this increased risk. For instance, it has been well established that females are at greater risk for autoimmune conditions due to an overactive and inflammatory immune system (26). Given that activated microglia and infiltrating monocytes are detectable in the early stages of AD (27), basal over-activation of innate immune cells in females could be a potential link to the increased prevalence of AD in women. Further studies have also shown that relative to healthy control females, females with AD possess a higher prevalence of less phagocytic pro-inflammatory macrophages (28). This could lead to lower Aβ uptake, and over-activation of surrounding immune cells. With this in mind, we determined that sex as a biological variable should be investigated in the present study, as sexually-dimorphic differences have been described for both immune function and risk for AD.

The aims of our study were two-fold. First, we aimed to study the effect of the *P522R* mutation on a peripheral macrophage population to determine how these cells may be differentially affected by the mutation. To do so, we assessed changes in major macrophage functions including phagocytosis and inflammatory cytokine secretion, as well as the transcriptional changes that underlie these functions. We utilized peritoneal macrophages (pMacs) which can be separated into two main populations, large peritoneal macrophages (LPMs) and small peritoneal macrophages (SPMs). In broad terms, LPMs are known to be resident to the peritoneal cavity, and SPMs are derived from blood-monocytes, which travel to the peritoneal cavity upon infection or other inflammatory events (29). Recent studies have shown that these recruited monocytes can, over time, differentiate into macrophages that closely mimic the resident LPMs (30). This population was selected over bone-marrow derived macrophages as peritoneal macrophages differentiate within the tissue, which allows them to mold to their environment and experience hormonal changes throughout the animal’s lifespan (29, 31, 32). Additionally, this population of peripheral macrophages is well characterized, lending to a more informed study that aims to assess tissue-resident macrophages specifically (29, 31, 32). Furthermore, the immune cells of the peritoneum are of great interest in relation to the gut, which has been identified as a peripheral reservoir for Aβ, and as a conduit of peripheral messengers to the brain via the gut-brain-axis (22, 33). Not only that, but immune responses occurring in this region have been shown to directly activate the brain’s insular cortex, a region known to be affected during the preclinical stages of AD (34), which can in turn store this immune-related information and alter the peripheral immune response if reactivated, further supporting a direct connection between the brain and peritoneum (35). Second, we aimed to address how sex may influence the effector functions of this population of macrophages, both with and without the *P522R* mutation, as knowing the baseline sex differences in wild-type (WT) mice would more easily allow us to assess additional effects of the *P522R* mutation. We found that the *P522R* mutation does indeed alter pMac function, supporting the idea that this mutation may potentially have protective effects on the organism due to its expression in the periphery and not just in brain-resident immune cells. However, we also found that the effects of this mutation are largely sexually dimorphic, indicating that future studies should utilize both male and female sexes to better inform potential drug development strategies for the clinic.

## METHODS

### Animals

Congenic mice expressing the *P522R* mutation in the *Plcg2* gene were purchased from the Jackson Laboratory (strain #029598) and housed at the University of Florida McKnight Brain Institute vivarium. Mice heterozygous for the *P522R* mutation were produced by breeding heterozygous mice and C57BL/6 control littermates. They were kept at 22°C in 60-70% humidity and on a 12-hour light-dark cycle and fed *ad libitum*. All animal procedures were approved by the University of Florida Institutional Animal Care and Use Committee and were in accordance with the National Institute of Health Guide for the Care and Use of Laboratory Animals (NIH Publications No. 80-23) revised 1996. Both male and female mice were bred and used in these experiments and were 3-5 months old at the time of sacrifice. Mice were sacrificed via cervical dislocation, and all experiments included 5-6 biological replicates.

### pMac isolation and culture

Three days prior to pMac extraction, mice were injected with 1mL of a 3% thioglycolate (Brewer) solution. Thirty minutes prior to thioglycolate injection mice were injected with 50uL of sustained-release Buprenorphine (ZooPharm) to alleviate pain. After three days, the mice were sacrificed via cervical dislocation. The abdomen was sprayed with 70% ethanol and the skin was split along the midline. Once the peritoneal cavity was exposed, a 10mL syringe filled with 10mL of cold RPMI (Gibco, 11875119) was injected into the peritoneal cavity using a 27G needle, avoiding blood vessels. Once the 10mL of cold RPMI was injected, another empty 10mL syringe with a 25-gauge needle was used to withdraw and place the peritoneal fluid into a 15mL Falcon collection tube. Any remaining fluid was aspirated via transfer pipette and placed into the collection tube. Once all mice were sacrificed and peritoneal fluid was collected into separate collection tubes, the fluid was passed through a 70uM nylon filter pre-wet with 5mL of cold HBSS-/- (Gibco, 14175103) into a 50mL conical tube. 5mL of cold HBSS-/- was then placed into the collection tube to rinse any remaining cells and passed through the filter as well. The filter was then washed with 5mL of HBSS-/- and the 50mL tube containing the fluid and HBSS-/- were centrifuged at 400*g* for 5 minutes at 4°C to pellet the cells. If any blood cells were present the pellet was resuspended in 1mL of ammonium-chloride-potassium lysis buffer for 1 minute and then centrifuged again at 400*g* for 5 minutes at 4°C. Once all cells were pelleted, the supernatant was aspirated and each pellet was resuspended in 3mL of plating media (RPMI, 10% fetal bovine serum, and 1% penicillin/streptomycin). Viable cells were identified and counted using trypan-blue exclusion on an automated cell counter (Countess™). The volume of growth media was adjusted so that cells were plated at a density appropriate for the intended assay (detailed below). Cells were incubated at 37°C, 5% CO_2_ for a minimum of 2-hours to allow macrophages to adhere. Wells were washed with sterile DPBS (Gibco, 14190235) to remove any non-adherent cells, and new, pre-warmed growth media was added.

### Flow Cytometry

For flow cytometry analysis, cells were plated at a density of 3×10^5^ cells/well in a 24-well plate and left to incubate overnight. 1 hour prior to collection, DQ-BSA green (Invitrogen, D12050) was added to each well at a concentration of 10ug/mL to measure protein degradation. After 1 hour, the cells were washed 3 times with sterile phosphate buffered saline (PBS) (Gibco, 10010023), detached from the plate via cell scraper, and then transferred to a 96-well v-bottom plate and centrifuged at 300*g* for 5 minutes at 4°C. The cells were then resuspended in 50uL of PBS containing fluorophore-conjugated antibodies for CD11b (1:100, Biolegend, 101216), MHCII (1:100, Biolegend, 107628), and Live/Dead stain (1:2000, Invitrogen, L34965). The cells were then incubated in the dark at 4°C for 20 minutes at which point they were centrifuged again at 300*g* for 5 minutes at 4°C. The cells were then washed with sterile PBS, and this process was repeated twice. The cells were then fixed in 50uL per well of 1% paraformaldehyde in PBS for 30 minutes in the dark at 4°C. After 30 minutes the cells were centrifuged at 300*g* for 5 minutes at 4°C, resuspended in 200uL of FACs buffer (PBS, 0.5 mM EDTA, 0.1% sodium azide), and taken to the Macs quant analyzer (Miltenyi) for flow cytometric analysis. A minimum of 100,000 events were captured per sample and data were analyzed using FlowJo version 10.6.2 software (BD Biosciences). When validating flow cytometry panels and antibodies, fluorescence minus one controls (FMOCs) were used to set gates and isotype controls were used to ensure antibody-specific binding.

### Phagocytosis Assay

For phagocytosis studies, cells were plated at a density of 5×10^4^ cells/well in a 96-well black wall plate (Cellvis, NC1648269) in triplicate. Cells were left to incubate for 6 hours at 37°C in a 5% CO_2_ humidified cell culture incubator. 2mL of Live Cell imaging solution (Invitrogen, A59688DJ) was added to each vial of pHrodo green *E. coli* bioparticles (Invitrogen, P35366), equating to a final concentration of 1mg/mL, and these solutions were sonicated for 10 minutes. After sonication, the media was removed from the cells in the plate and replaced with 100uL of *E. coli* bioparticle solution, at which point the plate was placed into an onstage incubator connected to an EVOS M7000 microscope to maintain the cells at 37°C and 5% CO_2_ humidified atmosphere throughout the duration of the live-cell imaging. GFP images were captured at 20X magnification with 5 images per well every 30 minutes for 15 hours within each well, and the EVOS Celleste (Version 5) software was used to count cells and determine the integrated optical density of GFP fluorescence within each well over time. Brightfield images were used for automated-focus throughout the time course. Treatment conditions were performed in triplicate wells and averaged for analysis.

### Multiplexed Immunoassays

For inflammatory cytokine secretion, isolated pMacs were plated at a density of 1×10^6^ cells/well in a 6-well plate and treated with either vehicle (PBS) or LPS (O111:B4 Sigma, L4391, Lot#0000185317) at a concentration of 10ng/mL. The cells were then placed in a humidified cell culture incubator at 37°C and 5% CO_2_. Conditioned media was collected at 2, 6, and 24 hours post stimulation for assessment of specific protein concentrations of TNF, IL-6, IL-1β, KC/GRO, IL-4, IL-10, IFN-γ, IL-5, IL-2, and IL-12p70 (MSD, K15048D-2) using multiplexed immunoassays on the MesoScale Discovery (MSD) platform. Samples were plated in duplicate at a 1:1 dilution with MSD diluent 41 and the rest of the experimental procedure was performed according to the manufacturer’s instructions. The plate was then analyzed on the MSD Quickplex machine and MSD software (Discovery Workbench Version 4.0). Cytokines with values below the lower limit of detection (LLOD listed below) were not quantified, these being 0.10pg/mL for TNF, 0.04pg/mL for IFN-γ, 0.11pg/mL for IL-1β, 0.22pg/mL for IL-2, 0.11pg/mL for IL-4, 0.06pg/mL for IL-5, 0.61pg/mL for IL-6, 0.94 pg/mL for IL-10, 9.95pg/mL for IL-12p70, and 0.24pg/mL for KC/GRO.

### RNA Isolation

For RNA extraction, cells were plated at 1×10^6^ cells/well in a 6-well plate and treated with either vehicle or LPS at 10ng/mL. The cells were placed back in a humidified cell culture incubator at 37°C and 5% CO_2_ for 6 hours. Buffer RLT from the RNeasy mini kit (Qiagen, 74104) was combined with β-mercaptoethanol (Sigma, 63689) at a concentration of 20uL of BME per 1mL of buffer RLT. Media was aspirated from the plate and then 350uL of RLT+BME solution was then added to each well to lyse the cells. Each well was scraped with a mini cell scraper (Fisher, NC0325221) and collected into a Qiashredder (Qiagen) column and centrifuged at 10,000*g* for 2 minutes. 350uL of 70% ethanol was added to the homogenized lysate and then transferred to a RNeasy spin column. The columns were then centrifuged for 30 seconds at 10,000*g* and afterwards the flow-through was discarded. 700uL of buffer RW1 was then added to each column and the columns were centrifuged at 10,000*g* for 30 seconds and the flow-through was discarded. 500ul of buffer RPE was added to each column and they were spun at 10,000*g* for 2 minutes and the flow-through was again discarded. This step was repeated twice, and the column was then placed into a 1.5mL collection tube, 30uL of elution buffer was added to the column, and then centrifuged for 1 minute at 10,000*g* to elute the RNA. Concentration of RNA was analyzed on a spectrophotometer, and RNA quality was assessed using Bioanalyzer 2100 (Agilent, Denmark).

### Gene expression analysis

For quantification of mRNA transcripts in pMacs, the NanoString nCounter platform and Mouse Immune Exhaustion Panel RNA (CAT#PSTD-M-EXHAUST-12) containing 785 genes was used. 25ng/ul of RNA was loaded and hybridized to the provided capture and reporter probes overnight at 65°C according to the manufacturer’s instructions. The samples were then added into the provided nCounter chip via the use of the NanoString MAX/FLEX robot. Unhybridized capture and reporter probes were removed, and the hybridized mRNAs were immobilized for transcript count quantification on the nCounter digital analyzer. The data was then imported into the nSolver analysis software v4.0 for quality check and analysis according to the manufacturer’s instructions. Raw counts were normalized to the geometric mean of the positive and negative controls and count expression was calculated via nSolver Advanced Analysis v2.0.134. Gene expression was considered significant if *p* < 0.05 after adjustment using the Benjamini-Hochberg method for false discovery rate, and if the fold change was greater than 1.5. Pathway analysis was performed using Enrichr for GO biological process analysis (36–38).

### Statistical Analysis

Statistical analysis was performed using Prism Graphpad version 10.2.2. Differences between groups was determined using either T-test with Welch’s correction or 2-way ANOVA with multiple comparisons using Tukey’s post-hoc correction. Results were considered statistically significant if *p* < 0.05. Each graph depicts individual data points with error bars representing the standard error of the mean (SEM).

## RESULTS

### The P522R mutation does not alter pMac sub-populations in male mice but increases antigen presentation

Before addressing cellular functionality, it is beneficial to first understand and characterize the macrophage populations present within our samples. As mentioned, pMacs are not a homogenous population, with SPMs arising from blood monocytes and LPMs being resident to the peritoneal cavity. To determine whether the *P522R* mutation has any effect on the frequencies of these cell populations we isolated pMacs from the peritoneal cavity of male mice, and then used flow cytometry to distinguish SPMs from LPMs on the basis of CD11b expression, as has been previously described (Supplemental Figure 1A) (39). Prior to isolation, we performed an intraperitoneal injection of a 3% thioglycolate solution. This step is crucial prior to isolation, as it allows us to harvest both LPMs and SPMs, not just the resident LPMs. However, we do allow a three-day rest period post injection and prior to cell isolation so that acute inflammation and cell activation induced by the injection subside (Figure 1).

**Figure 1:**
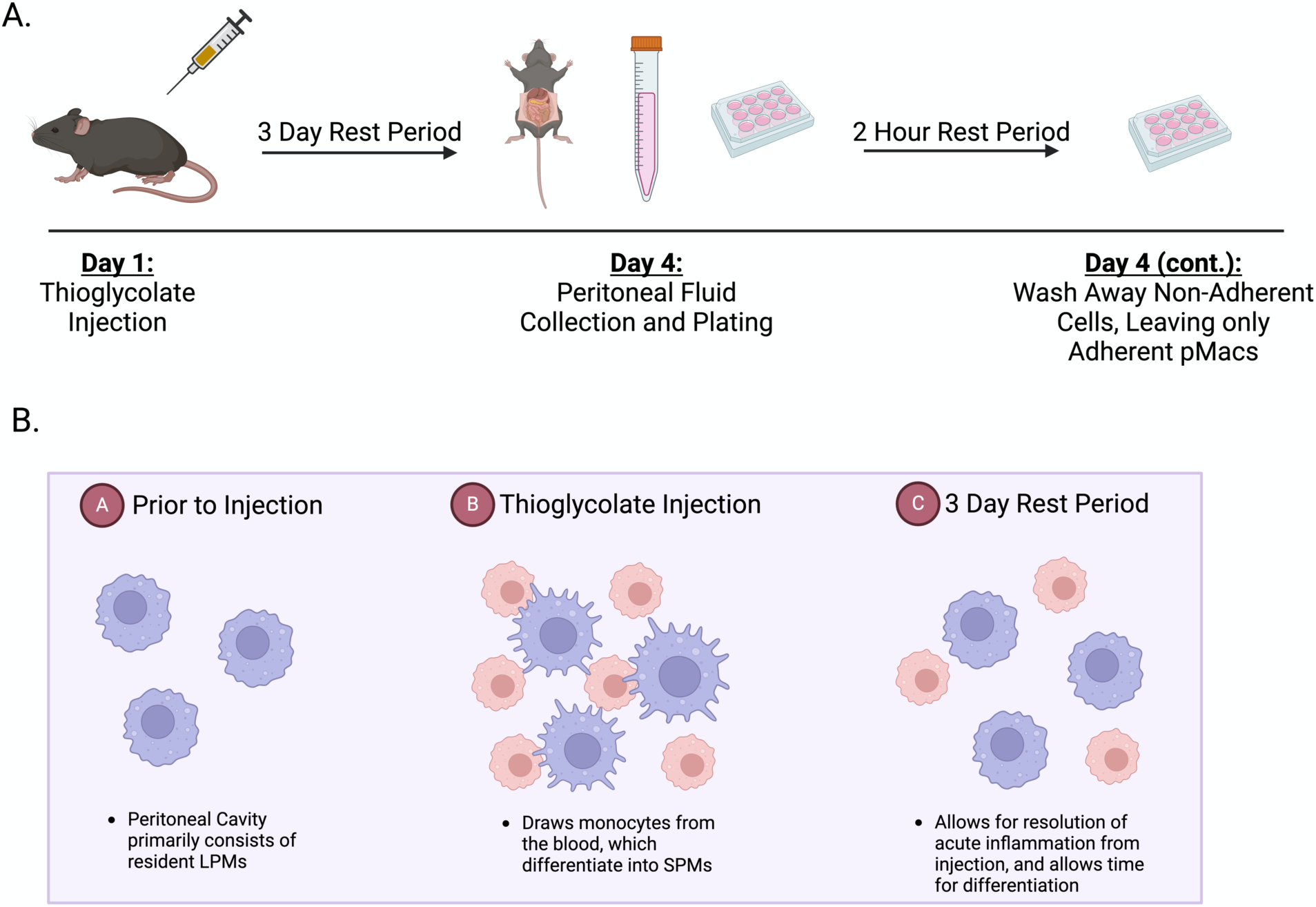
Schematic of pMac collection and the purpose of thioglycolate injections prior to cell isolation. Diagram describing the process of pMac isolation (A) and visual representation of the purpose of starting the process with a thioglycolate injection (B). Created with Biorender.com.

Additionally, after plating the cells collected from the peritoneum, we leave the cells to rest for two hours and then rinse with sterile DPBS and replace the media, as this allows only the adherent cells (pMacs) to remain and all other immune cell types to be washed away (Supplemental Figure 1B). We further confirmed LPM vs SPM populations by assessing canonical peritoneal macrophage markers of LPMs (CD80 and F4/80) and SPMs (Dectin1) (Supplemental Figure 1C-E). We found that male mice carrying the *P522R* mutation showed no difference in the frequency of either LPM or SPM sub-populations compared to WT male mice (Figure 2A, B). This was confirmed via raw counts (Supplemental Figure 2A-D). Next, we used major histocompatibility complex class II (MHCII) as a marker of antigen presentation. We found that although LPM and SPM populations were not affected by the *P522R* mutation in male mice, antigen presentation on LPMs was. A higher frequency of LPMs were MHCII+ in *P522R* males compared to LPMs from WT males (Figure 2C).

**Figure 2:**
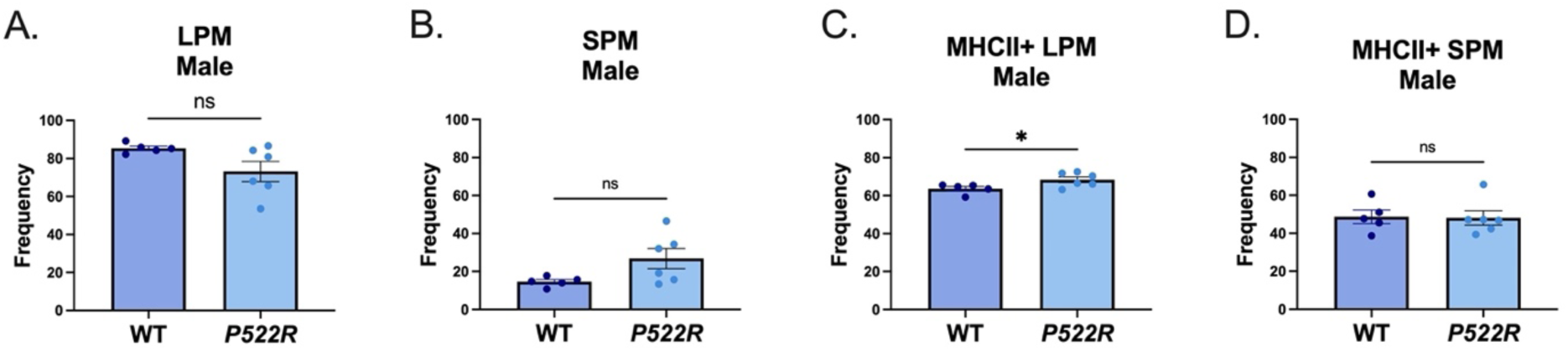
The *P522R* mutation does not alter pMac sub-populations in male mice but increases antigen presentation. To determine the prevalence of different sub-populations of macrophages in our *ex vivo* cultures pMacs from male WT and *P522R* animals were isolated, plated, and non-adherent cells were washed away after a 2-hour rest period and populations of (A) LPM (B) SPM (C) MHCII+ LPM and (D) MHCII+ SPM were quantified via flow cytometry. Data was quantified via T-test with Welch’s correction. Groups are considered non-significant if *p*>0.05 and significantly different if *p*<0.05.

We did not find any significant differences between *P522R* and WT males in MHCII expression on SPMs (Figure 2D). Representative plots for this analysis can be found in Supplemental Figure 2E and 2F. These data indicate that in males the *P522R* mutation does not affect the makeup of peritoneal macrophage sub-populations, but does increase the amount of resident LPMs displaying antigens.

### The P522R mutation alters both sub-populations and antigen presentation in pMacs from female mice

Based on the literature indicating distinct sex-differences in pMac populations, we aimed to determine whether the *P522R* mutation would have similar effects on population frequencies and antigen presentation in female mice as was seen in male mice (40). However, we found that the *P522R* mutation had quite different effects in female mice compared to those seen in male mice. Female mice carrying the *P522R* mutation exhibited significant increases in the frequency of LPMs, and significantly lower frequencies of SPMs compared to WT females (Figure 3A, B).

**Figure 3:**
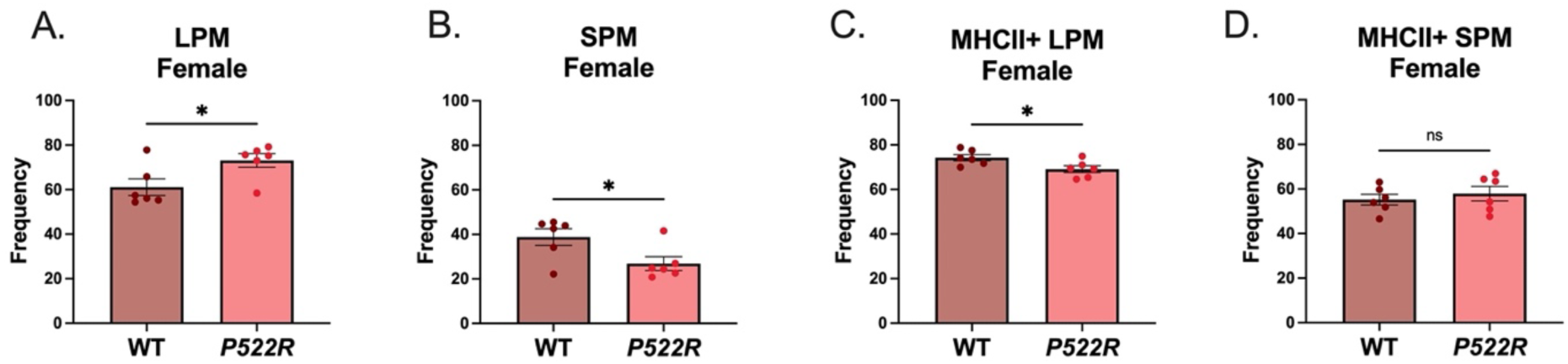
The *P522R* mutation alters both sub-populations and antigen presentation in pMacs from female mice. To determine the prevalence of different sub-populations of macrophages in our *ex vivo* cultures pMacs from female WT and *P522R* animals were isolated, plated, and non-adherent cells were washed away after a 2-hour rest period and populations of (A) LPM (B) SPM (C) MHCII+ LPM and (D) MHCII+ SPM were quantified via flow cytometry. Data was quantified via T-test with Welch’s correction. Groups are considered non-significant if p>0.05 and significantly different if *p*<0.05.

After reviewing the counts, we determined that these effects were primarily driven by an increase in the raw count of LPMs in females that carry the *P522R* mutation (Supplemental Figure 3A-D). Furthermore, contrary to the effect seen in pMacs from males, the *P522R* mutation’s presence in females actually decreased the frequency of MHCII+ LPMs compared to pMacs from WT females (Figure 3C). But similar to males, there were no observable differences between the frequencies of MHCII+ SPMs between WT or *P522R* females (Figure 3D), representative plots for which can be found in Supplemental Figure 3E and 3F.

These data support a role for the *P522R* mutation in altering macrophage sub-populations within the peritoneum of female mice. However, these effects are largely sex-specific, and even bi-directional in the case of antigen presentation. These data begin to confirm a role for the *P522R* mutation in peripheral tissue resident macrophages, and further support previously published data indicating significant sex-differences in immune cell populations and function.

### The P522R mutation reduces phagocytic capacity and lysosomal protease activity in pMacs from males

After determining that the *P522R* mutation can alter pMac populations in female mice, and antigen presentation on LPMs in a bidirectional manner in females and males, we aimed to determine whether this mutation also has functional consequences in pMacs from each sex. Macrophages are known to have many dynamic functions, yet phagocytosis and cytokine secretion are among the most recognized, both of which PLCG2 is actively involved in. We began with assessing phagocytosis capabilities in pMacs from male WT and *P522R* mice, as the ability to uptake and degrade bacterial/viral particles is among the most important functions of macrophages in the fight against infection. Furthermore, this function has been directly associated with AD, as *E. coli* particles and LPS particles have been shown to associate directly with pathology in the brains of AD patients (41). Additionally, AD progression has been correlated with infection, emphasizing the importance of the pMacs ability to uptake and degrade bacterial particles (42). To assess the phagocytic capabilities of these macrophages, we utilized fluorescently labeled *E. coli* bioparticles which only fluoresce after being internalized and exposed to the low pH associated with the lysosome, thereby reporting on the extent of bacterial uptake by the cells. Interestingly, we found that pMacs from *P522R* male mice performed less uptake of the fluorescent *E. coli* over the course of 15 hours than did pMacs from WT male mice (Figure 4A-C). Up until approximately the 6 hour mark, the *P522R* and WT pMacs performed very similar amounts of *E. coli* uptake, but the *P522R* pMacs were not able to keep up with the WT pMacs, which seemingly performed better (Figure 4B). Although the macrophages were able to take up the *E. coli* particles, we further assessed whether pMacs from males displayed any differences in lysosomal function, or the ability to successfully break down that which has been phagocytosed. To do so, we treated the cells with DQ-BSA, a substrate for proteases that only fluoresces after it has been cleaved within the lysosome, and quantified via flow cytometry.

**Figure 4:**
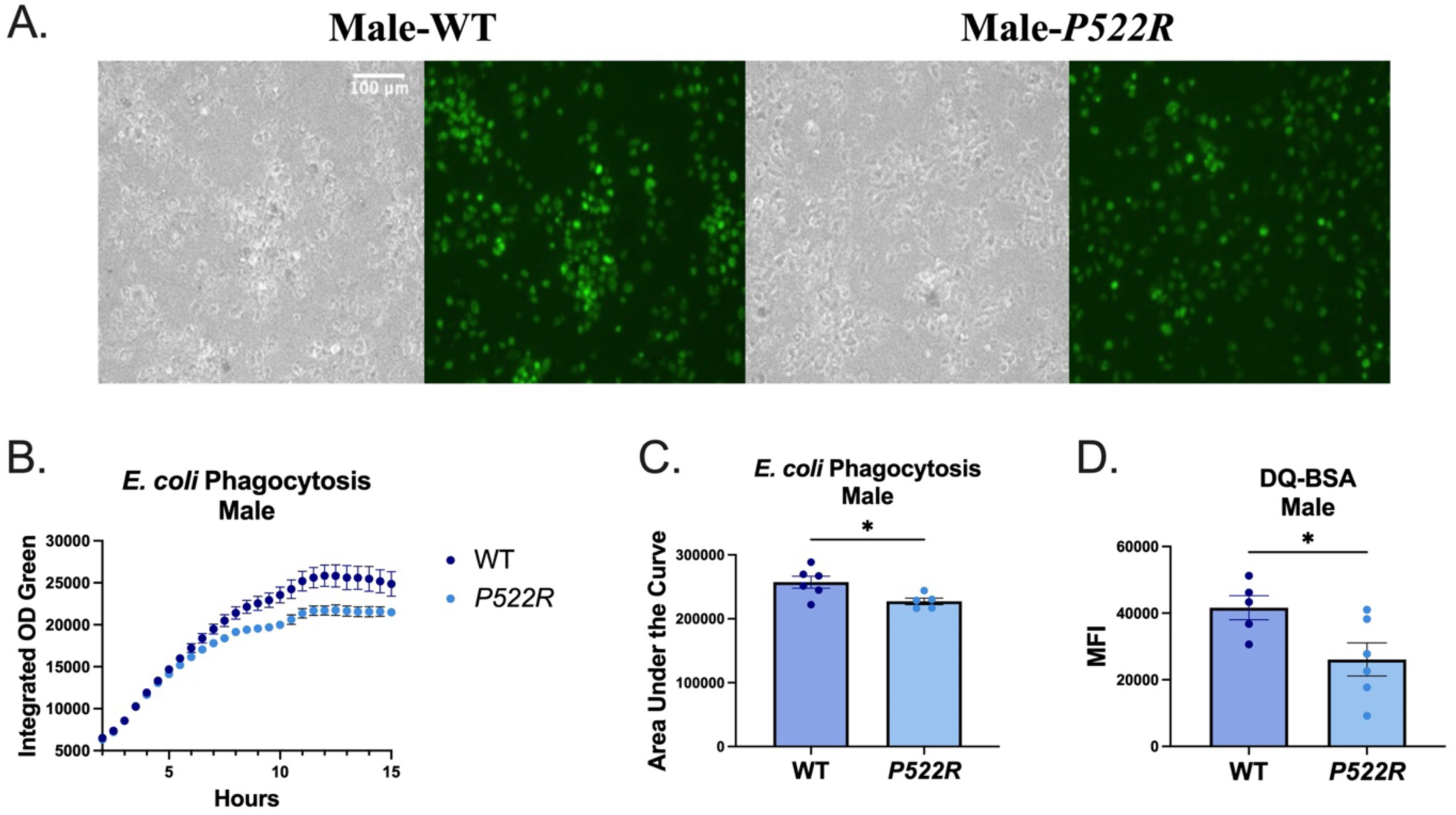
The *P522R* mutation reduces phagocytic capacity and lysosomal protease activity in pMacs from males. pMacs from WT and *P522R* males were plated and incubated for 6 hours, at which point 200ug of *E. coli* was added to each experimental well to assess *E. coli* uptake. (A) Representative brightfield images and GFP fluorescence for each group are presented at 20X magnification. (B) Integrated optical density of GFP fluorescence was plotted over 15 hours. (C) Groups were compared statistically via area under the curve. 1 hour prior to cell collection DQ-BSA was added to the plate and then cells were collected for flow cytometry and (D) DQ-BSA MFI is represented. Groups are considered non-significant if *p*>0.05 and significantly different if *p*<0.05.

Again, we found that pMacs from WT males performed better than pMacs from *P522R* males, which exhibited lower lysosomal protease activity compared to pMacs from WT males (Figure 4D). These data indicate that the *P522R* mutation does indeed alter phagocytosis and lysosomal activity in males, but these alterations may not be beneficial to pMac function.

### The P522R mutation does not alter phagocytic capacity in pMacs from females, but increases lysosomal protease activity

As the *P522R* mutation has been associated with protection, the decrease in phagocytic capacity and lysosomal protease activity seen in pMacs from *P522R* males was quite surprising. We repeated both the *E. coli* uptake assay, as well as the DQ-BSA lysosomal protease activity assay in pMacs from WT and *P522R* females to determine whether similar patterns were seen in females. Similar to what we found in our assessment of pMac populations, the *P522R* mutation had different effects in pMacs from females compared to pMacs from males. Although there appeared to be a visual difference in GFP fluorescence between pMacs from *P522R* and WT females, quantification showed that uptake of *E. coli* in pMacs from females was largely unaffected by the *P522R* mutation (Figure 5A-C). However, lysosomal protease activity measured via fluorescence of the DQ-BSA probe was shown to be increased in pMacs from *P522R* females (Figure 5D).

**Figure 5:**
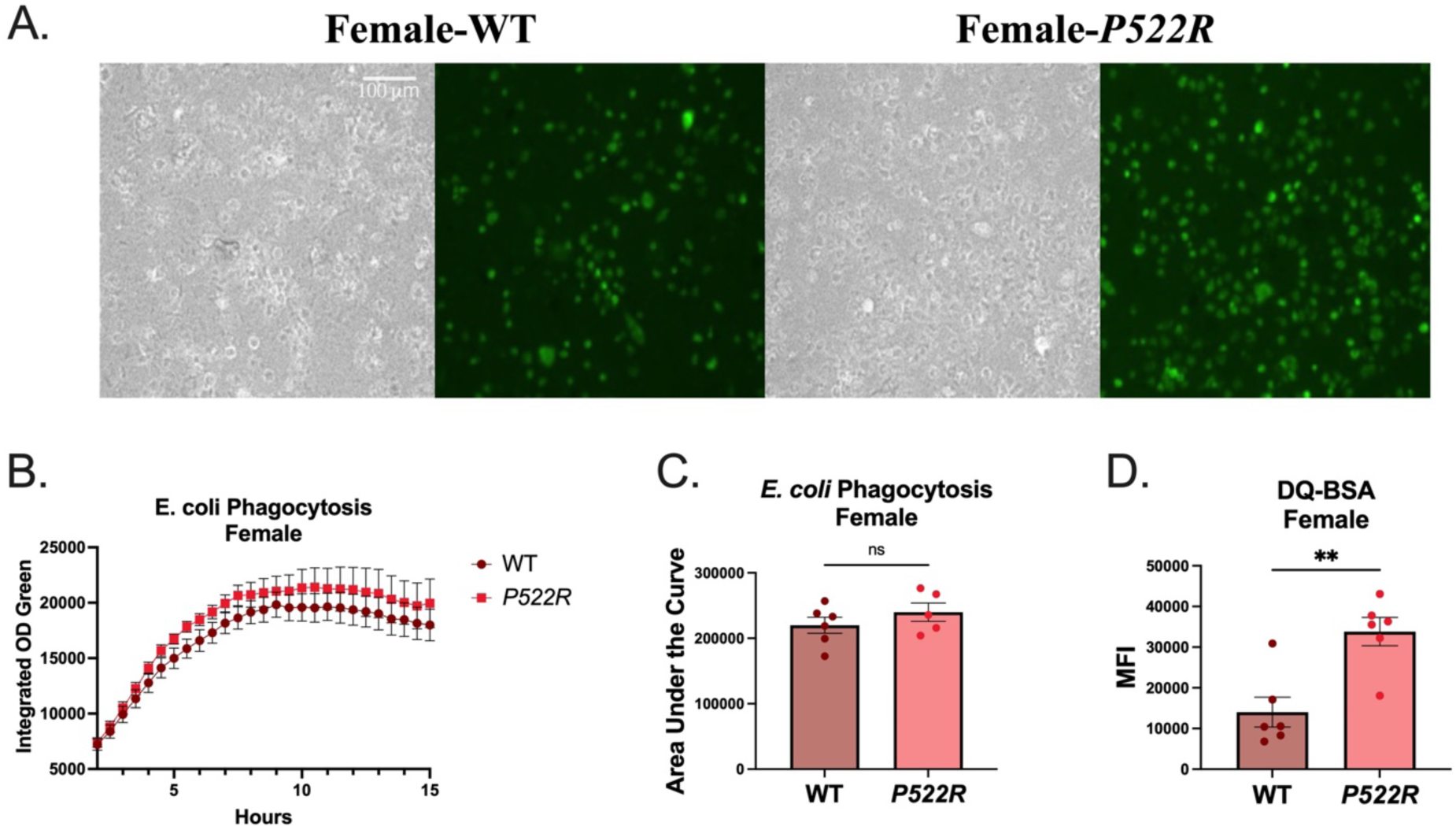
The *P522R* mutation does not alter phagocytic capacity in pMacs from females, but increases lysosomal protease activity. pMacs from WT and *P522R* females were plated and incubated for 6 hours, at which point 200ug of *E. coli* was added to each experimental well to assess *E. coli* uptake. (A) Representative brightfield images and GFP fluorescence for each group are presented at 20X magnification. (B) Integrated optical density of GFP fluorescence was plotted over 15 hours. (C) Groups were compared statistically via area under the curve. 1 hour prior to cell collection DQ-BSA was added to the plate and then cells were collected for flow cytometry and (D) DQ-BSA MFI is represented. Groups are considered non-significant if *p*>0.05 and significantly different if *p*<0.05.

These data indicate that the *P522R* mutation may confer functional advantages in lysosomal activity in pMacs from females, but not in males.

### The P522R mutation modestly increases inflammatory cytokine secretion by pMacs from males

As mentioned, the other function of macrophages that has been linked to PLCG2 function is cytokine secretion. We have already demonstrated that the *P522R* mutation has sex-dependent effects on phagocytosis and lysosomal function, and we hypothesized that it would also alter cytokine secretion by pMacs, likely also in a sex-dependent manner. To test this, we treated pMacs from WT and *P522R* males with LPS and measured the protein concentration of cytokines within the media at 2, 6, and 24 hours post-LPS treatment. We found that vehicle treated cells produced very low concentrations of inflammatory cytokines, with no difference between WT and *P522R* cells, further supporting that our *ex vivo* cultures are not activated or inflammatory at baseline conditions (Figure 6A-H). For this reason, we treated with the immune cell stimulant LPS to elicit an inflammatory response, as although there were no genotype differences at baseline, the different genotypes may respond differently to inflammatory stimuli.

**Figure 6:**
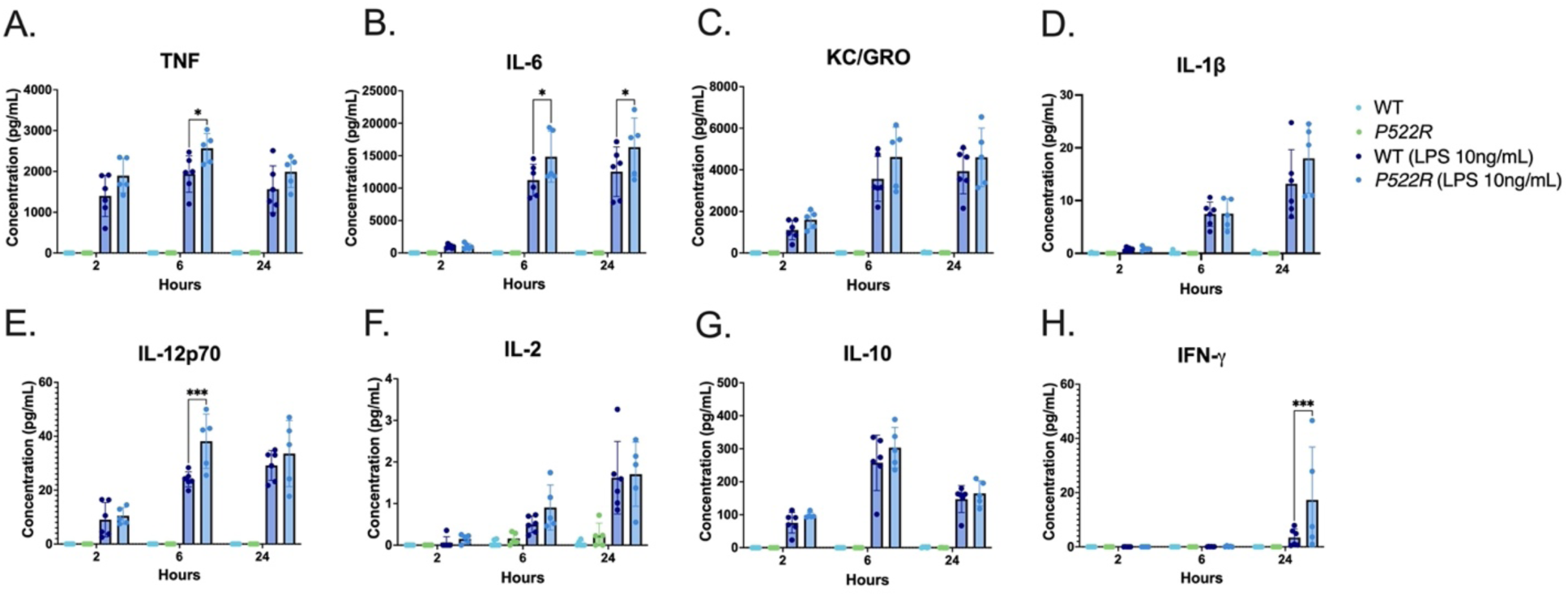
The *P522R* mutation modestly increases inflammatory cytokine secretion by pMacs from males. pMacs from male WT and *P522R* mice were plated and treated with LPS, and their conditioned media was collected at 2, 6, and 24 hours post LPS stimulation. Concentrations of (A) TNF (B) IL-6 (C) KC/GRO (D) IL-1β (E) IL-12p70 (F) IL-2 (G) IL-10 and (H) IFN-γ were then assessed via MesoScale Discovery immunoassays. Data were quantified via two-way ANOVA using Tukey’s post-hoc correction for multiple comparisons. Groups are considered non-significant if *p*>0.05 and significantly different if *p*<0.05.

Indeed, we found that upon LPS stimulation, pMacs from *P522R* males showed modest increases in inflammatory cytokine secretion of TNF, IL-6, and IL-12p70 at 6 hours post-LPS stimulation (Figure 6A, B and E), and IFN-γ at 24 hours post-LPS stimulation compared to pMacs from WT males (Figure 6H). We found no differences between genotypes in secretion of KC/GRO, IL-1β, IL-2, or IL-10 even after LPS stimulation (Figure 6C, D, F and G). These data support that the *P522R* mutation has a modest effect on pMac cytokine secretion in males.

### The P522R mutation increases inflammatory cytokine secretion by pMacs from females

To determine whether these same stimulation dependent effects were seen in females, we repeated this same experiment but with pMacs from WT and *P522R* females. Similar to pMacs from males, we found the pMacs from both WT and *P522R* females produce very low levels of inflammatory cytokines when not stimulated with LPS (Figure 7A-H), supporting that, like our male cultures, the female *ex vivo* cultures are not inflammatory or activated prior to stimulation. Similar to what we saw in our male data, LPS stimulation did reveal genotype-dependent differences in cytokine secretion, however these differences were different and more prevalent than in our male cultures. For example, at 6 hours post-LPS stimulation, pMacs from *P522R* females were secreting more TNF, KC/GRO, IL-1β, and IL-12p70 than pMacs from WT females (Figure 7A, C, D, and E). At 24 hours post-LPS stimulation, pMacs from *P522R* females were secreting more TNF, IL-6, KC/GRO, IL-1β, and IL-2 compared to pMacs from WT females (Figure 7A-D, and F). We found no genotype dependent differences in IL-10 or IFN-γ at any time point (Figure 7G, H). So, although the *P522R* mutation caused increases in cytokine secretion by pMacs from both males and females, the pMacs from females seemed to be more affected by the presence of the mutation and produced significantly more inflammatory cytokines over time.

**Figure 7:**
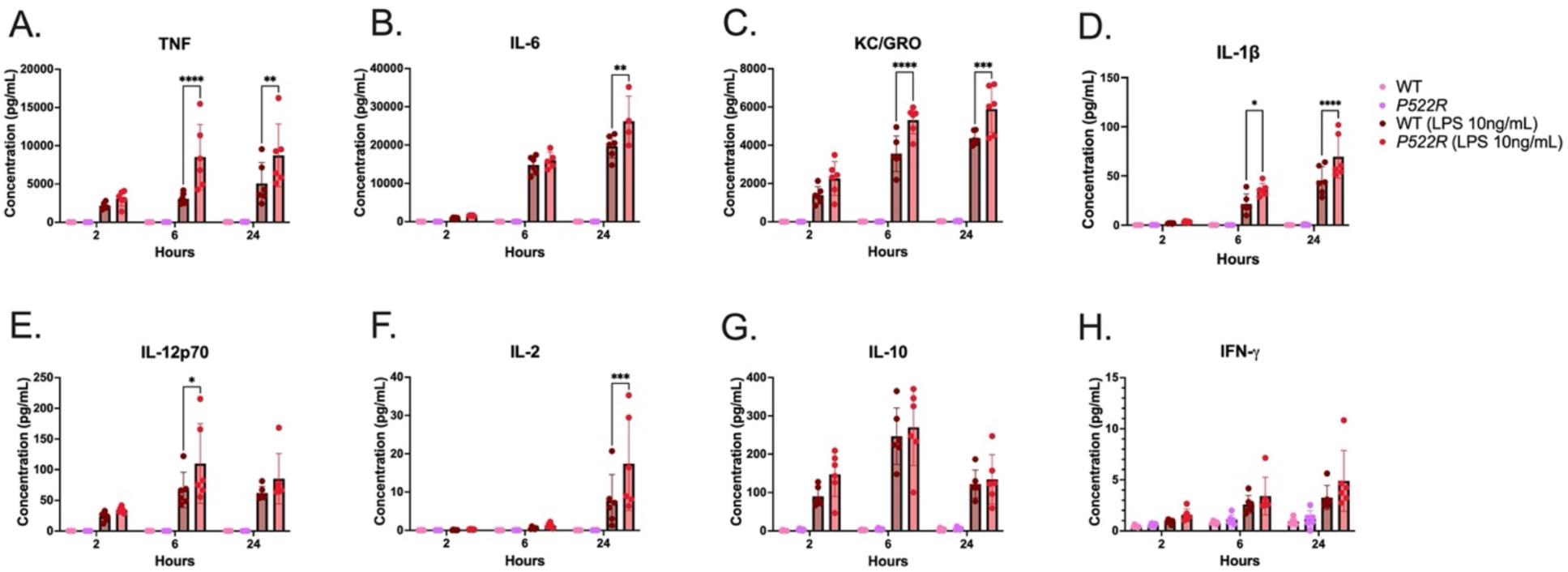
The *P522R* mutation increases inflammatory cytokine secretion by pMacs from females. pMacs from **fe**male WT and *P522R* mice were plated and treated with LPS, and their conditioned media was collected at 2, 6, and 24 hours post-LPS stimulation. Concentrations of (A) TNF (B) IL-6 (C) KC/GRO (D) IL-1β (E) IL-12p70 (F) IL-2 (G) IL-10 and (H) IFN-γ were then assessed via MesoScale Discovery immunoassays. Data were quantified via two-way ANOVA using Tukey’s post-hoc correction for multiple comparisons. Groups are considered non-significant if *p*>0.05 and significantly different if *p*<0.05.

### Genes selectively upregulated in P522R males are associated with negative regulation of the immune response

In our selected assays, the *P522R* mutation seemed to have a dampening effect on pMacs from males, indicating a potential sex-specific role for the mutation. Furthermore, many of these differences were more noticeable after treatment with some sort of immune cell stimulant, such as *E. coli* or LPS, supporting the idea that the *P522R* mutation’s effects may be activation-dependent, and not as prevalent under baseline conditions. To attempt to identify molecular pathways associated with the effects of stimulation, we performed a targeted transcriptomic analysis using Nanostring nCounter to assess 785 genes associated with immune cell function in pMacs from both WT and *P522R* males. We did so both at baseline and after treatment with LPS to determine whether these effects were truly activation dependent. As expected, we did not find any significant differences in gene expression between pMacs from WT and *P522R* males under baseline conditions (Supplemental Figure 5A). However, after treatment with LPS, differences between the genotypes did begin to arise.

Volcano plots representing each genotype’s response to LPS stimulation are represented in Figure 8A and 8B, with 8A showing pMacs from WT male’s response to LPS, and 8B showing pMacs from *P522R* male’s response to LPS. We found that many of the differentially expressed genes (DEGs) induced by LPS stimulation were shared between pMacs from WT and *P522R* males, encompassing 264 shared DEGs (Figure 8C). However, there were a number of genes that were genotype specific, including 13 DEGs specific to *P522R* males, and 21 specific to WT males (Figure 8C). We took each of these sets of genotype specific genes and assessed what pathways were associated with them. We found that the DEGs specific to pMacs from WT males were associated with pathways including calcineurin-NFAT signaling cascade, regulation of calcium ion transport across the plasma membrane, calcineurin-mediated signaling, and protein modification/phosphorylation (Figure 8D). Many of these pathways involve calcium signaling, and the calcineurin cascade which in macrophages has been shown to down-regulate TLR mediated activation pathways (43). Contrarily, the DEGs specific to pMacs from *P522R* males were associated with pathways including negative regulation of immune responses, regulation of type II interferon production, regulation of NK T cell activation, and response to UV (Figure 8E). These results suggest that the *P522R* mutation induces gene expression that has a dampening effect on pMac immune responses in males.

**Figure 8:**
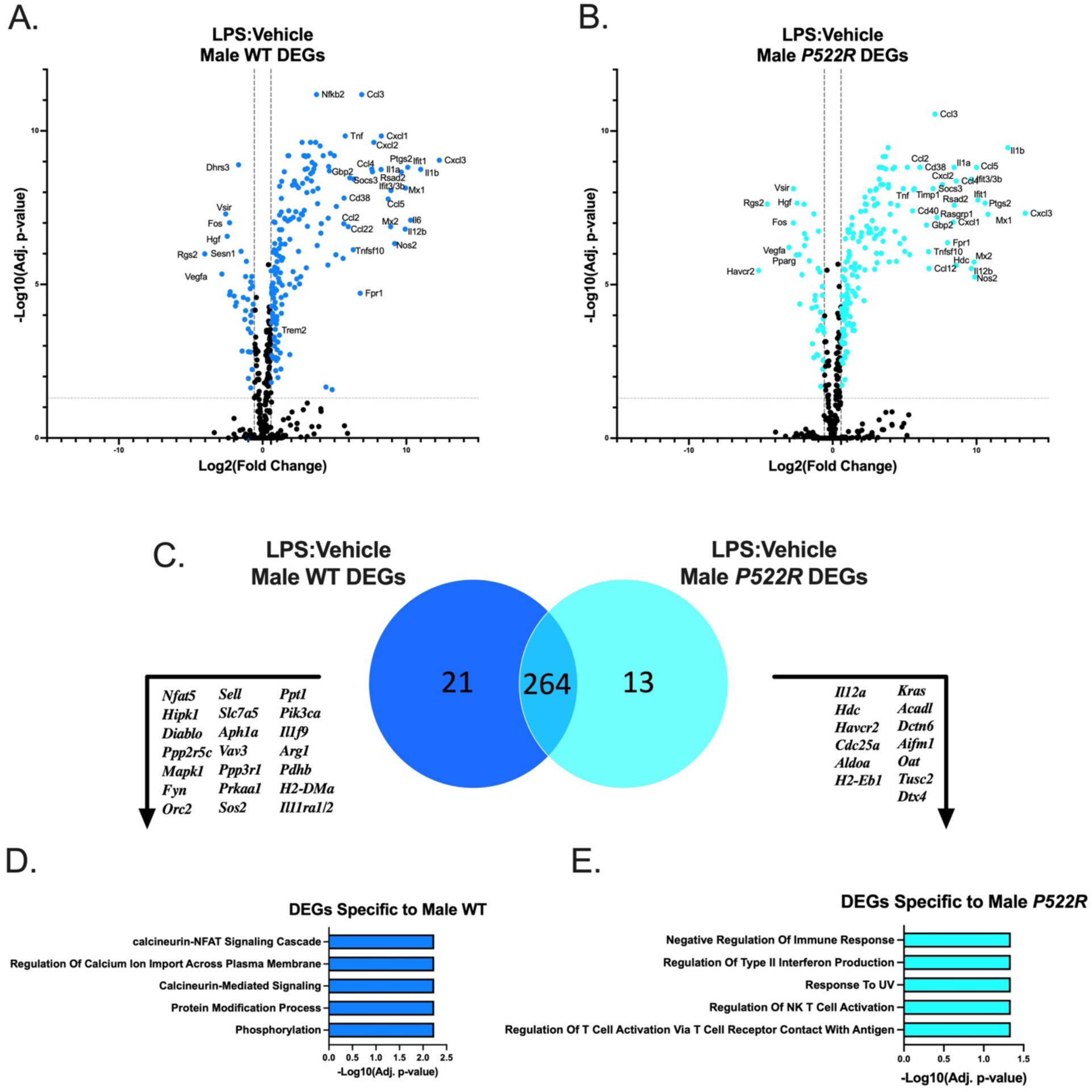
Genes selectively upregulated in *P522R* males are associated with negative regulation of the immune response. pMacs from male WT and *P522R* mice were plated and treated with either vehicle or LPS (10ng/mL) and left to incubate for 6 hours. Cells were then collected and RNA was isolated for gene expression analysis. (A) Volcano plot comparing gene expression of LPS treated pMacs from male WT mice to gene expression of vehicle treated pMacs from male WT mice, horizontal grey dashed line indicating adjusted *p*<0.05, and vertical dashed lines indicating fold change of greater than 1.5. (B) Volcano plot comparing gene expression of LPS treated pMacs from male *P522R* mice to gene expression of vehicle treated pMacs from male *P522R* mice, horizontal grey dashed line indicating adjusted *p*<0.05, and vertical dashed lines indicating fold change of greater than 1.5. (C) Venn diagram comparing number of DEGs between pMacs from WT males and pMacs from *P522R* males after LPS stimulation. (D) GO biological process pathways associated with DEGs specific to pMacs from WT males in response to LPS stimulation. (D) GO biological process pathways associated with DEGs specific to pMacs from *P522R* males in response to LPS stimulation.

### Genes selectively upregulated in P522R females are associated with cytokine signaling and apoptosis

As so far we have seen sex-dependent differences in pMac responses to the *P522R* mutation, we did targeted transcriptomics on our female samples as well to determine which pathways might be differentially implicated in the female’s varied responses. Similar to our male data, when pMacs from females are not treated with LPS, there are no significant gene expression differences between pMacs from WT females and *P522R* females (Supplemental Figure 5B).

However, upon LPS treatment, different responses to this activating stimulus occur. DEGs are represented in volcano plots for pMacs from WT females after LPS treatment (Figure 9A), and pMacs from *P522R* females after LPS treatment (Figure 9B). Many of the DEGs implicated in the cellular response to LPS are the same between pMacs from WT and *P522R* females, comprising 190 shared DEGs (Figure 9C), but each genotype did again have specific DEGs related to them, with 50 DEGs specific to WT females, and 12 DEGs specific to *P522R* females in response to LPS. These distinct gene groups were once again analyzed via GO biological process pathway analysis, and the results were, unsurprisingly, quite distinct from the male groups. We found that DEGs specific to pMacs from WT females were associated with pathways including regulation of defense response, regulation of immune response, regulation of autophagy, and negative regulation of the inflammatory response (Figure 9D). DEGs specific to pMacs from *P522R* females on the other hand were associated with pathways including cytokine mediated signaling, apoptotic processing, extrinsic apoptotic signaling, and positive regulation of peptidyl-tyrosine phosphorylation (Figure 9E). This suggests that the *P522R* mutation induces genes related to the upregulation of immune responses in pMacs from females.

**Figure 9:**
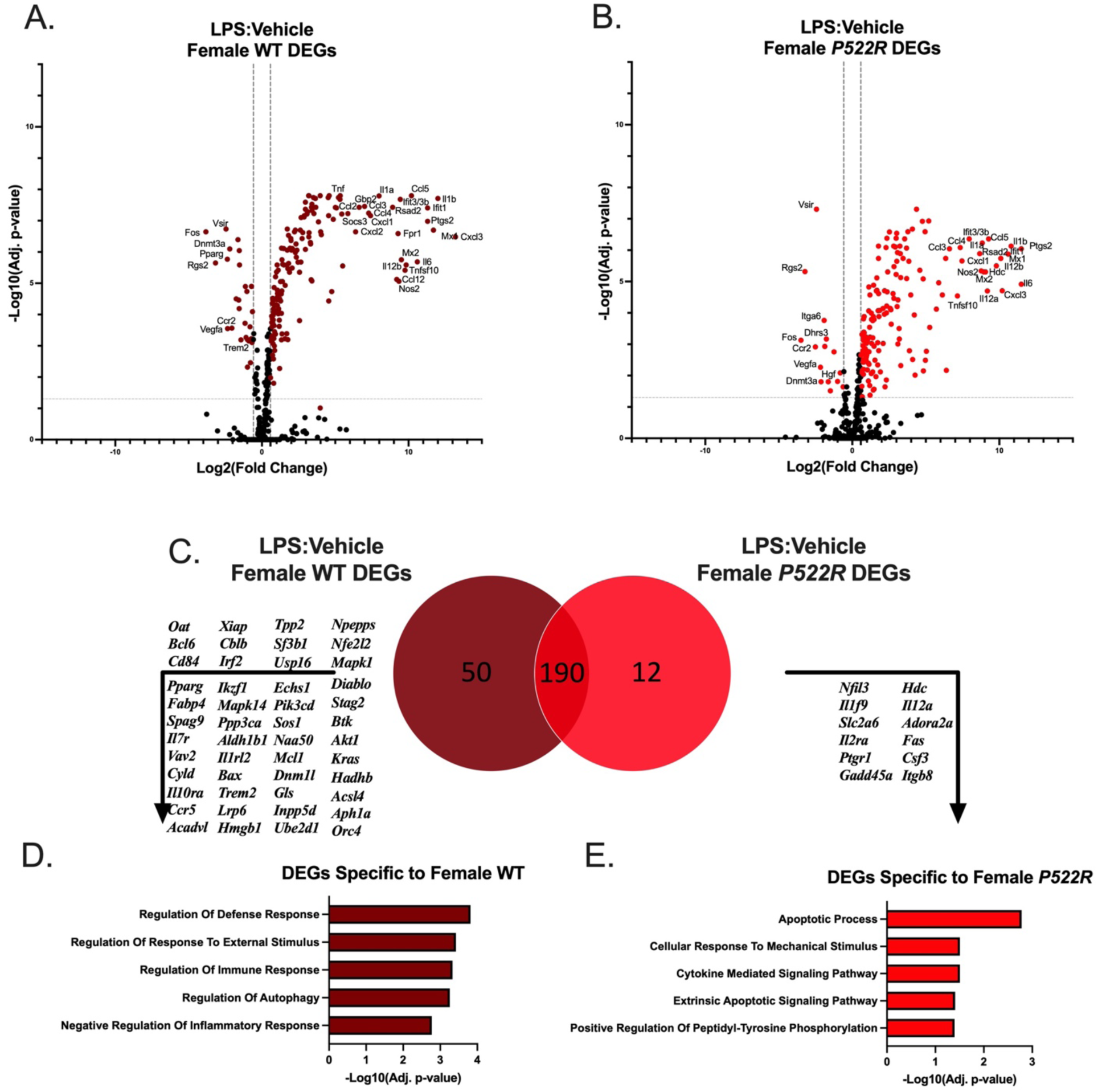
Genes selectively upregulated in *P522R* females are associated with cytokine signaling and apoptosis. pMacs from female WT and *P522R* mice were plated and treated with either vehicle or LPS (10ng/mL) and left to incubate for 6 hours. Cells were then collected and RNA was isolated for gene expression analysis. (A) Volcano plot comparing gene expression of LPS treated pMacs from female WT mice to gene expression of vehicle treated pMacs from female WT mice, horizontal grey dashed line indicating adjusted *p*<0.05, and vertical dashed lines indicating fold change of greater than 1.5. (B) Volcano plot comparing gene expression of LPS treated pMacs from female *P522R* mice to gene expression of vehicle treated pMacs from female *P522R* mice, horizontal grey dashed line indicating adjusted *p*<0.05, and vertical dashed lines indicating fold change of greater than 1.5. (C) Venn diagram comparing number of DEGs between pMacs from WT females and pMacs from *P522R* females after LPS stimulation. (D) GO biological process pathways associated with DEGs specific to pMacs from WT females in response to LPS stimulation. (D) GO biological process pathways associated with DEGs specific to pMacs from *P522R* females in response to LPS stimulation.

## DISCUSSION

Numerous studies have implicated the immune system as a player in the progression of neurodegenerative disorders, both within the brain and in the periphery. This includes evidence that peripheral monocytes and tissue-resident macrophages participate in Aβ plaque uptake, both around the brain and in peripheral tissue reservoirs (22–24). Furthermore, macrophages are known to be one of the main producers of inflammatory cytokines, which, when chronic, can exacerbate neurodegeneration (2–4), but when acute, act as messengers to brain immune cells and signals to other local immune cells (44–46). Therefore, it is of critical importance to investigate the nature of peripheral immune mechanisms to identify potential new targets for therapeutic intervention. With this objective in mind, many groups have studied the effects of the *Plcg2-P522R* protective mutation in myeloid cells, primarily microglia, to try to determine how this protection is being conferred since the effects manifest themselves in improved cognitive function. However, the effect of the *P522R* mutation on peripheral tissue-resident macrophages, which have become of increasing importance in the context of AD, had yet to be determined. Additionally, we assessed how the mutation’s effects may differ between sexes, as it is well known that both AD risk and immune system function is highly affected by sex. Herein, we show that the *P522R* mutation does indeed confer functional alterations in the periphery, but these changes seem to vary by sex.

When assessing pMac sub-populations, we found that the *P522R* mutation had different effects that were dependent on sex. In WT male mice, we found that our *ex vivo* cultures were predominantly comprised of LPMs, making up approximately 80% of the population (the other portion being made up of SPMs), and we found this to be the same in cultures obtained from *P522R* males, indicating that the mutation does not have an effect on pMac sub-populations in male mice. In cultures obtained from female WT mice however, we found that only around 60-65% of the population on average was comprised of LPMs. The addition of the *P522R* mutation altered this significantly, bringing this percentage up to closer to 80% LPMs. These data indicate that the mutation alters pMac sub-populations in females but not males, and interestingly seemed to bring female sub-population composition up to meet that which was found in males. Although the canonical sub-populations known as LPMs and SPMs are recognized as separate groups (29), recent studies have shown that not only can SPMs continue to alter their functional and transcriptional profiles to look more like LPMs, and even occupy that same niche over time, but they actually do so more readily in males (30, 40). It might be possible then that the *P522R* mutation somehow ablates this male-bias, making it possible for the infiltrating cells to become more LPM-like in females over time as is seen in males. However, it could also stand to reason that this increase in LPMs in *P522R* females may happen prior to/separate from our interventions, potentially during the time of peritoneum colonization by the early resident macrophages. Our data is not enough from which to draw substantiated conclusions on this topic, however it can be concluded that the *P522R* mutation does alter sub-populations (particularly LPMs) in female mice via a currently unknown mechanism. Based on our data and the aforementioned studies assessing the dynamic properties of these macrophages to shift from one sub-group to another at any particular time, we decided to perform the rest of the experiments on the entire population of pMacs available within a well, instead of trying to isolate LPMs and SPMs specifically for functional assays. Although possible, the ability of these cells to take on characteristics of the other group make this unnecessary, as these sub-populations are not static regardless. We do however, use our flow data pertaining to the makeup of our populations in each sex to influence our interpretation of functional results, but we do not rely solely on this for the aforementioned reasons.

The first functions we assessed were phagocytosis and lysosomal activity, these being the ability to take up particles, and the ability to break them down, respectively. The literature addressing the effect of the *P522R* mutation on phagocytosis is inconsistent, showing both increased and decreased phagocytosis (14–16). This discrepancy was hypothesized to be due to the size of the cargo, with studies showing lower phagocytosis of larger particles such as zymosan and *E. coli*, and more phagocytosis of smaller-sized dextrans (14, 15). We show here that this observed difference in phagocytosis caused by the *P522R* mutation could also be due to the sex in which the testing was performed. In our cultures, we found that the *P522R* mutation did alter phagocytosis and lysosomal activity in pMacs, but in opposing directions in males and females. Our data showed that pMacs from *P522R* males exhibited less uptake of *E. coli* particles over time compared to pMacs from WT males. In females, these same differences were not found, and the *P522R* mutation seemingly had little effect on *E. coli* uptake over time. We further observed changes in lysosomal activity, with the *P522R* mutation significantly reducing lysosomal protease activity in males, and significantly increasing it in females, further supporting the sex-dependent differences this mutation confers. It should be noted however, that differences in phagocytosis in males were not observed until around the 6-7 hour mark, meaning that initial uptake of *E. coli* was similar between groups, but the pMacs from *P522R* males may become exhausted earlier or more rapidly. Taken together, these results suggested the potential of the *P522R* mutation to be positive in females, but potentially negative in males via the reduction in bacterial uptake and degradation.

Further functional assays surveyed inflammatory cytokine secretion by pMacs from females and males, as previous studies showed that the *P522R* mutation was capable of increasing cytokine secretion in microglia (13). Our data indicated that this effect is also present in pMacs, but to different extents in males and females. We measured cytokine release both at baseline and post-LPS to determine whether there may be differences prior to any sort of external stimulation. In both male and female cultures, we saw that at baseline, regardless of genotype, the pMacs were not producing inflammatory cytokines, supporting that our *ex vivo* culture model allows for the cells to return to baseline non-inflammatory conditions. For this reason, we chose to treat the cells with the immune cell stimulant LPS to elicit an inflammatory response that could be compared between the groups. After LPS stimulation, pMacs from *P522R* males showed increased release of TNF, IL-6, IL-12p70, and IFN-γ over 24 hours compared to pMacs from WT males, while pMacs from *P522R* females showed increased release of TNF, IL-6, KC/GRO, IL-1β, IL-12p70, and IL-2 over 24 hours compared to pMacs from WT females. Interestingly, the *P522R* mutation had no effect on the release of IL-10, a canonical cytokine involved in the resolution of inflammatory responses, in either females or males. Although it could be assumed that these increases in inflammatory cytokines could be negative, our study only looked at acute cytokine secretion, which is likely to have been an adaptive and beneficial response.

In our assays assessing the effects of the *P522R* mutation on pMac function and population dynamics we found different functional outcomes depending on sex. Furthermore, we found that many of these differences were more/only prominent after some form of cell activation either via *E. coli* or LPS treatment. This led us to believe that the *P522R* mutation may exert stronger effects upon cell activation in females and males, and we wondered whether we might be able to potentially determine some cellular pathways implicated in these differential responses, and therefore we performed targeted transcriptomics to assess immune cell specific gene expression changes. First, we compared gene expression in pMacs from *P522R* males to pMacs from WT males and found no differences at baseline. From that point, we compared gene expression after LPS stimulation between these two groups to determine what changes might be occurring due to the *P522R* mutation in response to cellular activation. What we found was quite fascinating. We saw that in response to LPS stimulation, the DEGs specific to *P522R* males were related to pathways associated with negative regulation of the immune response. This lines up with the functional data that revealed that in pMacs from males, the *P522R* mutation decreases phagocytosis and lysosomal activity, and does not exert as strong of an effect on inflammatory cytokine secretion as is seen in females. In females however, we found that the DEGs specific to pMacs from *P522R* females were quite different, being related to cytokine mediated signaling pathways, positive regulation of peptidyl-tyrosine phosphorylation, and apoptotic signaling. Tyrosine phosphorylation has been characterized as a main event following LPS stimulation in macrophages which aids in cellular activation (47), in line with the observed increase in lysosomal activity and cytokine secretion in females. Furthermore, cytokine mediated pathways and apoptotic signaling can go hand in hand, as cytokines are main promoters of apoptosis, particularly TNF (48, 49). It appears based on our transcriptomic data that the *P522R* mutation activates different intracellular signaling pathways and concurrent gene expression in response to LPS leading to the different outcomes in response to activation seen between male and females.

At this time, we cannot say directly how these sex-dependent responses may contribute to the neuroprotection associated with this mutation. However, we can say with certainty that peripheral immune cells, particularly peritoneal macrophages, are affected by the presence of this mutation, and show transcriptional, lysosomal, phagocytic, and inflammatory alterations due to its presence, particularly after an inflammatory challenge. With recent studies indicating a role for peripheral macrophages in the progression of AD, this study opens up new avenues of interest for more in-depth studies of the contribution of peripheral macrophages to the *P522R* mutation’s protective effects. It should be noted however, that these responses and functional differences may not solely be due to intrinsic changes in macrophages due to the *P522R* mutation. As we isolated these cells from the peritoneum, a location filled with many immune cell types (many of which would also carry the KI *P522R* mutation), some of these functional changes could be due to the macrophages’ direct interaction with/response to other mutated immune cells which might leave lasting impacts on macrophage function. While we hypothesize that the functional changes seen in this study are due to direct intrinsic changes in macrophages brought about by the *P522R* mutation, which is supported by the studies showing intrinsic mechanisms of the *P522R* mutation in similar cell type microglia (12, 13), we cannot completely rule out the contribution of the peritoneal environment prior to our isolation protocol. Additionally, we provide ample evidence to necessitate the use of both sexes in any future study aiming to assess the protective effects of this mutation. Many drugs aiming to cure or mediate disease have been pulled from the market due to unforeseen issues arising in one sex or another (50, 51). Over recent years the issue of sex-differences in biomedical research has come to the forefront, and we provide further evidence to support the advantages of utilizing both sexes in pre-clinical studies whenever possible. Although this study does not provide an exact mechanism by which these peripheral macrophages may contribute to the protective effects of this mutation, it provides compelling reasons to support future inquiries into these questions, which we aim to do moving forward using *in vivo* models of neurodegeneration to determine whether these same functional alterations directly impact neuroinflammation and cognitive performance. It further supports the importance of including sex as a biological variable in studies as well, showing that not only can the magnitude of response to protective genetic mutations be different between sexes, but also the directionality of response can vary by sex and will have implications for future therapeutic interventions.

## CONCLUSION

Inflammation and the cells of the peripheral immune system have been implicated in the progression of AD. Therefore, identifying pathways that could be targeted in therapeutic interventions will give the field the potential to modulate these dysregulated immune cell functions. Others have demonstrated that the *P522R* mutation has the ability to enhance important immune functions in microglia, such as phagocytosis and cytokine secretion. Herein we provide the first evidence that the *P522R* mutation also modulates peripheral tissue-resident macrophage functions, and that these effects are highly sex-dependent. Taken together, these results reveal that peripheral immune cell changes should be taken into account when assessing the mechanism(s) by which deleterious/beneficial genetic mutations confer their effects, as microglia may not be the only cells contributing to the functional outcome of the genetic variant, and that sex as a biological variable should be an integral part of any study looking to assess peripheral inflammatory mechanisms that are likely to affect brain function.

## LIST OF ABBREVIATIONS

PLCG2: phospholipase C gamma 2
AD: Alzheimer’s disease
Aβ: amyloid-beta
TREM2: triggering receptor expressed on myeloid cells 2
TLR: toll-like receptor
PIP2: phosphatidylinositol (4,5) bisphosphate
DAG: diacylglycerol
IP3: inositol 1,4,5 triphosphate
LPM: large peritoneal macrophage
SPM: small peritoneal macrophage
WT: wild-type
PBS: phosphate buffered saline
FMOC: fluorescence minus one control
MSD: MesoScale Discovery
LLOD: lower limit of detection
SEM: standard error of the mean
MHCII: major histocompatibility complex II
LPS: lipopolysaccharide
DEG: differentially expressed gene

## DECLARATIONS

### Ethics approval and consent to participate

Mice were utilized in accordance with protocols approved by the Institutional Animal Care and Use Committee of the University of Florida.

### Consent for publication

Not applicable.

### Availability of data and materials

All data are included in the present study or contained within the supplemental files. Raw data can be provided upon request.

### Authors Contributions

HAS and MGT conceived the study. HAS designed the experiments, performed them, and analyzed and interpreted the data. MLB, RLW, JJ, KBM, CC, NN, and AMT assisted with various experimental procedures. HAS designed all figures. HAS and MGT wrote and edited the manuscript. All authors approved the final version of the manuscript and figures.

## Acknowledgements

The authors thank the Tansey, Khoshbouei, and Chakrabarty labs for useful discussions.

