## Supplemental Figures for "Alzheimer’s disease-associated protective variant Plcg2-P522R modulates peripheral macrophage function in a sex-dimorphic manner"

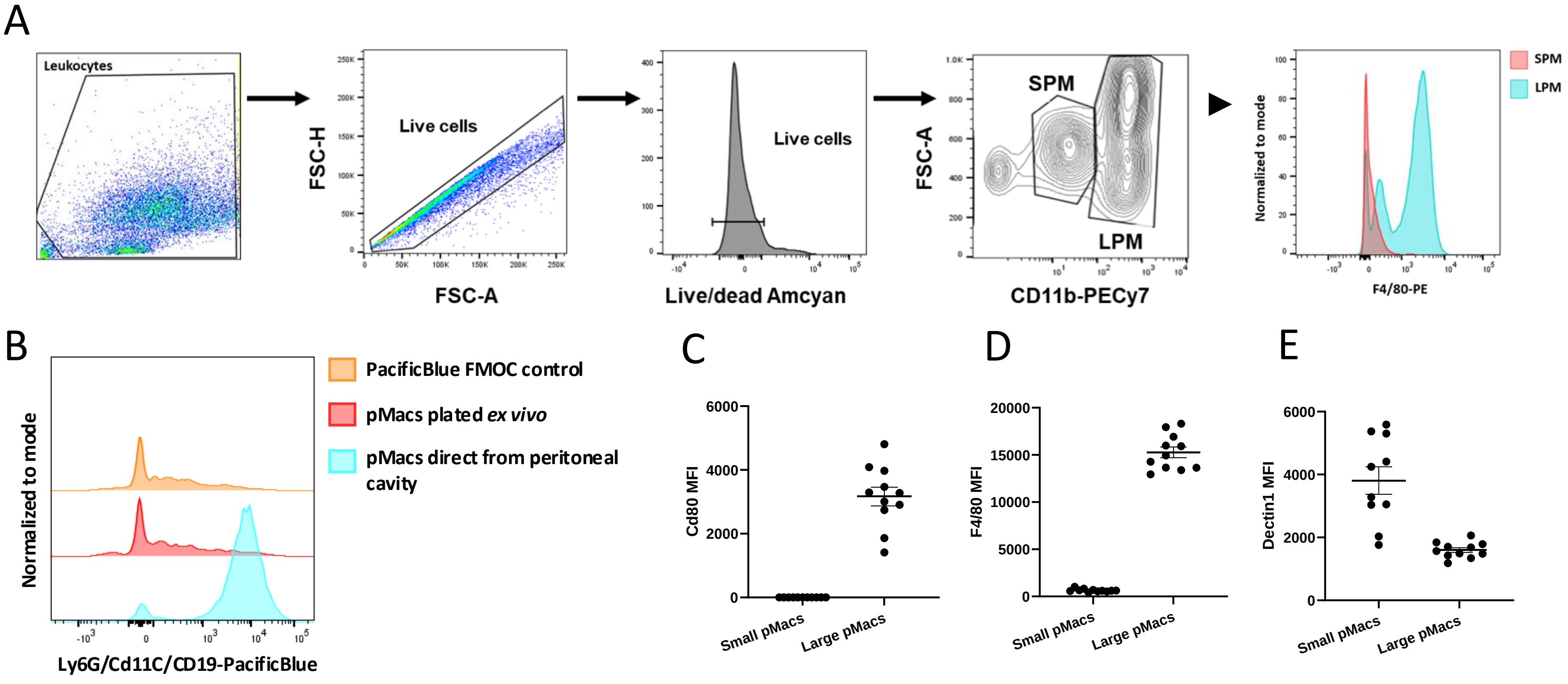


**Supplementary Figure 1:** ***Flow cytometry gating strategy used to identify LPM and SPM populations.*** (A) pMac populations were gated on a basis of CD11b expression, with LPMs expressing high levels of CD11b, and SPMs exhibiting low levels of CD11b. This gating was performed on live cell populations. (B) Flow cytometry plot showing the difference in populations between fluid directly from the peritoneal cavity, and cells after plating and washing away excess non-adherent cells. SPM and LPM populations were further confirmed by assessing canonical peritoneal macrophage markers (C) CD80 MFI (D) F4/80 MFI and (E) Dectin1 MFI.

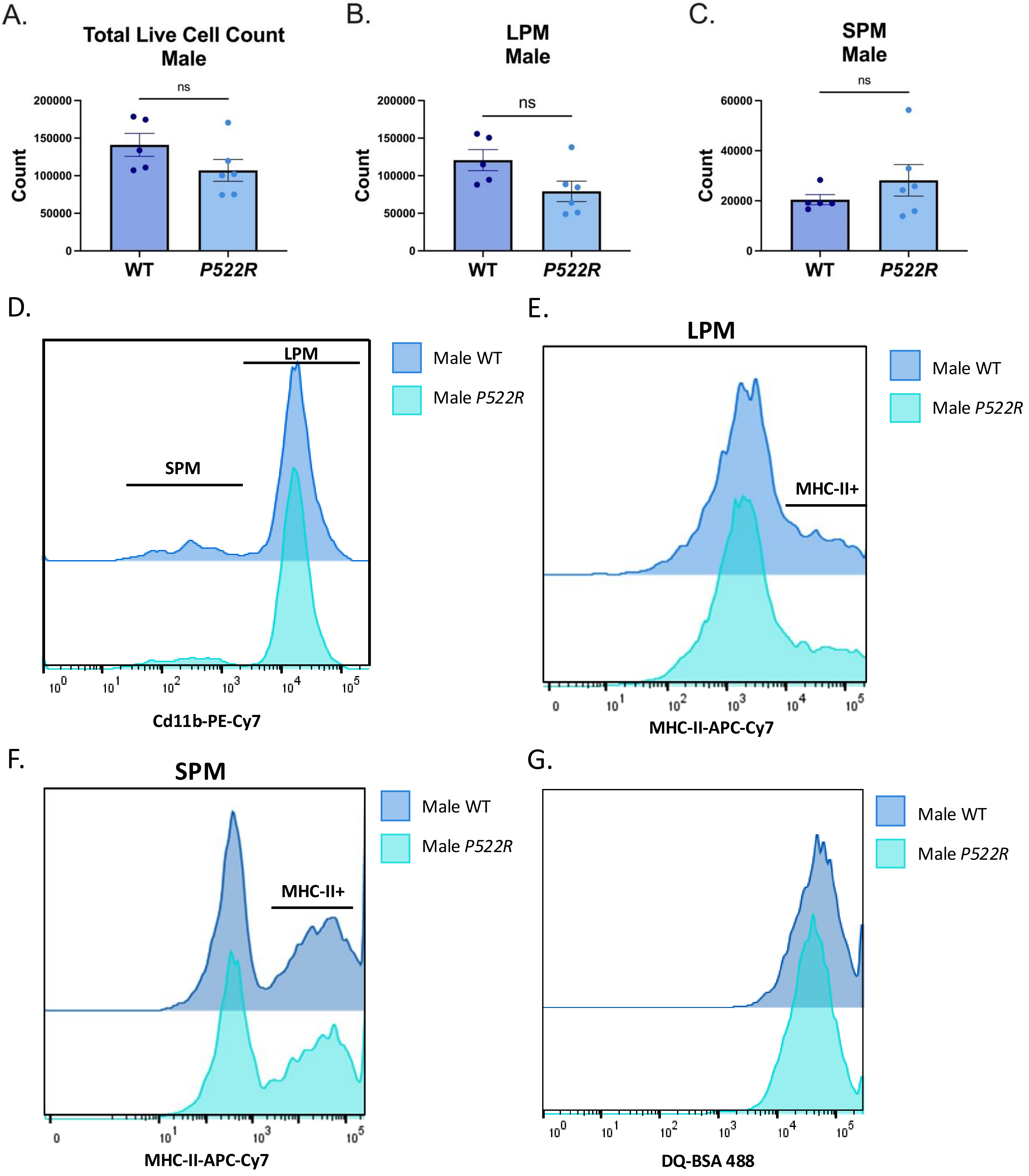


**Supplementary Figure 2: Raw counts and FACs plots for male pMac flow analysis.** pMacs from WT and *P522R* males were plated, incubated for 18 hours, and then taken for flow cytometry to assess sub-populations. Total live cells counts (A), LPM counts (B), and SPM counts (C) are represented for each genotype. FACs plots representing LPM vs SPM (D), MHCII on LPMs (E), MHCII on SPMs (F), and DQ-BSA (G) for each group are also given.

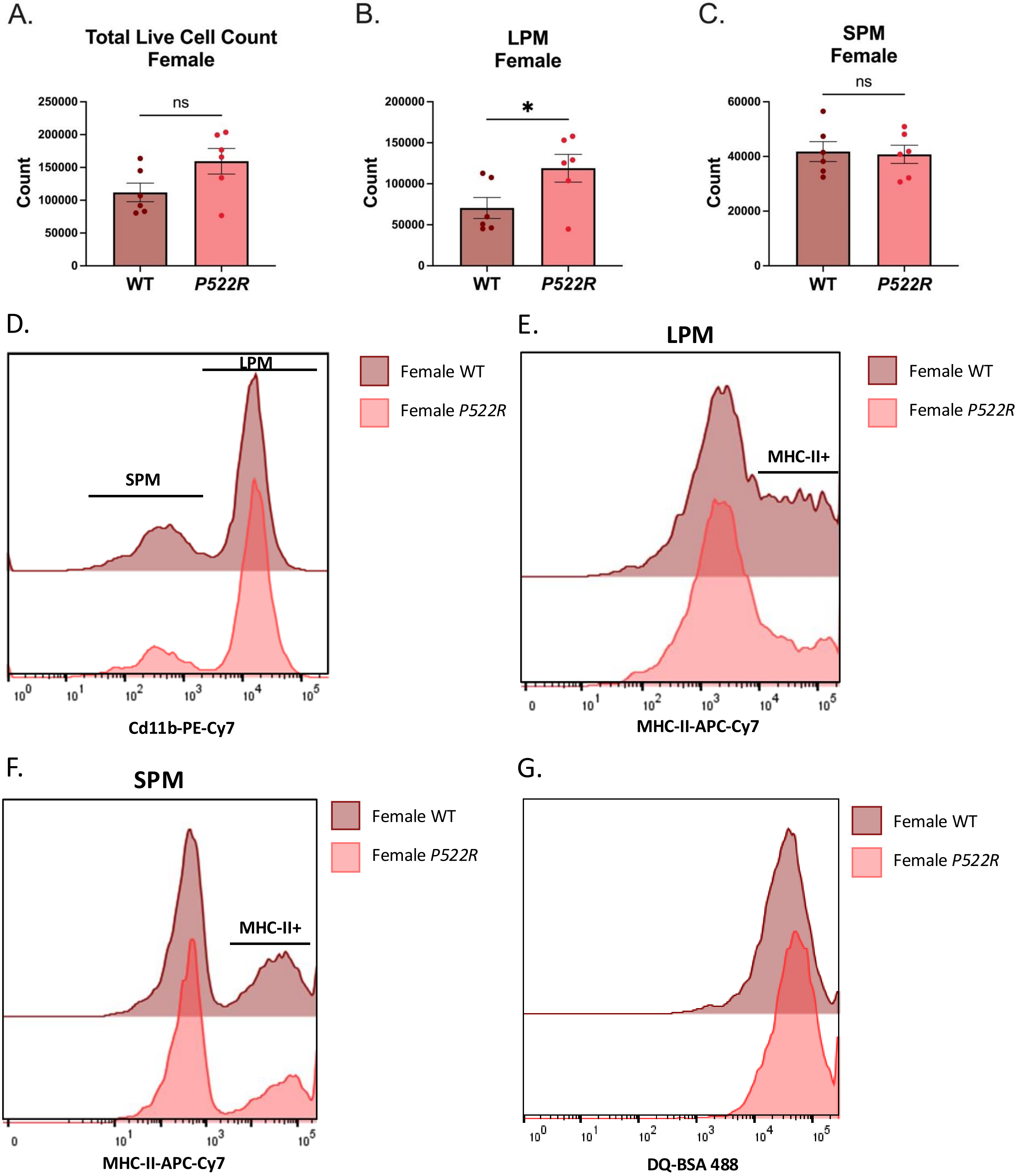


**Supplementary Figure 3: Raw counts and FACs plots for female pMac flow analysis.** pMacs from WT and *P522R* females were plated, incubated for 18 hours, and then taken for flow cytometry to assess sub-populations. Total live cells counts (A), LPM counts (B), and SPM counts (C) are represented. For each genotype. FACs plots representing LPM vs SPM (D), MHCII on LPMs (E), MHCII on SPMs (F), and DQ-BSA (G) for each group are also given.

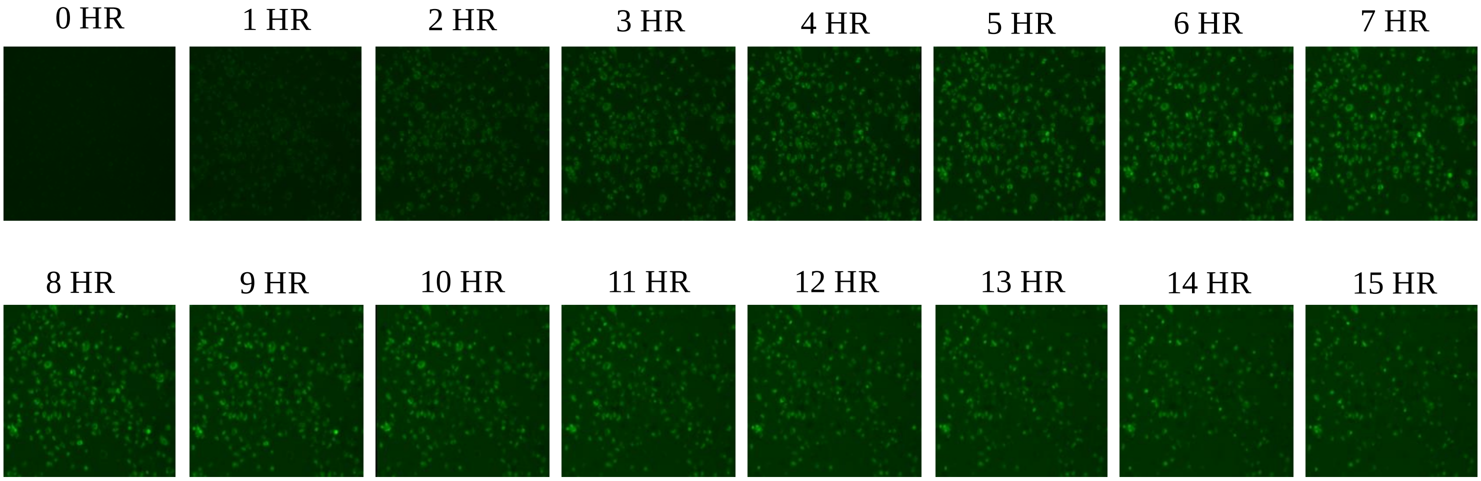


**Supplementary Figure 4: Representative time series images of phagocytosis assay GFP fluorescence across 15 hours.**

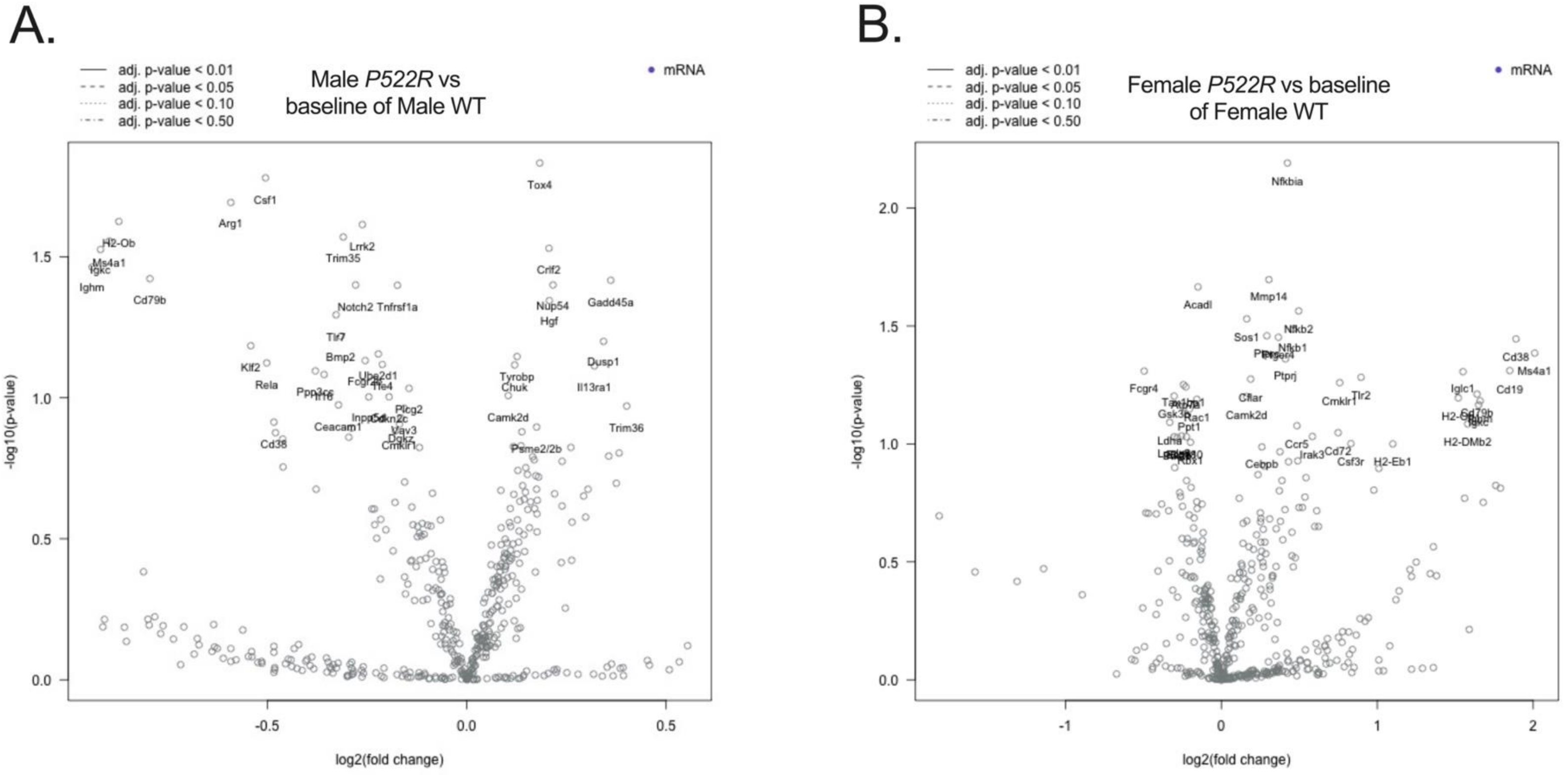


**Supplementary Figure 5: Gene expression volcano plots comparing the *P522R* mutation's effects within sex with no LPS stimulation.** Gene expression from pMacs from *P522R* males compared to pMacs from WT males (A), and gene expression from pMacs from *P522R* males compared to pMacs from WT males (B).
